# Machine learning-guided reconstruction of cytoskeleton network from Live-cell AFM Images

**DOI:** 10.1101/2024.03.21.584818

**Authors:** Hanqiu Ju, Henrik Skibbe, Masaya Fukui, Shige H. Yoshimura, Honda Naoki

## Abstract

How actin filaments (F-actins) are dynamically reorganized in motile cells at the level of individual filaments is an open question. To this end, we developed a high-speed atomic force microscopy (HS-AFM) to live-imagine intracellular dynamics of the individual F-actins. However, noise and low resolution made it difficult to fully recognize individual F-actins in the HS-AFM images. To tackle this problem, we developed a new machine learning method that quantitatively recognizes individual F-actins. The method estimates F-actin orientation from the image while improving the resolution. We found that F-actins were oriented at ±35° toward the membrane in the lamellipodia, which is consistent with Arp2/3 complex-induced branching. Furthermore, in the cell cortex our results showed non-random orientation at four specific angles, suggesting a new mechanism for F-actin organization demonstrating the potential of our newly developed method to fundamentally improve our understanding of the structural dynamics of F-actin networks.

## Introduction

Actin cytoskeletons are important for a wide range of cellular processes, such as morphogenesis and motility, due in part to their ability to facilitate polymerization/depolymerization, branching, and crosslinking between filaments^1–3^. Actin reorganization is regulated by intracellular signaling via actin-associated molecules. Many studies have live-imaged the activity of signaling molecules such as Rho GTPases and examined the relationship between molecular activities and morphological changes^4,5^. Our understanding of the relationship between the molecular activities and morphological changes of the cytoskeleton, however, is currently limited. Indeed, it is unclear how signaling molecules regulate the polymerization/depolymerization, branching, and crosslinking of each F-actin, and subsequently, how this affects morphological changes. To fill this knowledge gap, each F-actin in living cells should be tracked.

Cytoskeleton architecture has predominantly been observed using electron microscopy (EM) at a high spatial resolution^6,7^. F-actins can assemble into several formations, including branched networks in the lamellipodia, bundled filaments in the filopodia, and actomyosin in stress fibers^1,8–11^. F-actin architecture was recently analyzed quantitatively in terms of branching point density, actin filament density and the angular distribution of actin filaments from EM images^6,10,12^. However, EM observations are limited to dead, rather than living, cells. It has consequently been impossible, until date, to observe the dynamics of each F-actin in living cells using EM.

Tip-scan high-speed atomic force microscopy (HS-AFM) imaging systems have been established for live-cell imaging of cellular surfaces^13,14^. We have developed a new HS-AFM that can help to visualize the structural dynamics of the cortical actin network^15^. Using this technology, the polymerization rate and frequency of filament synthesis were measured in living cells. However, precisely recognizing the F-actin architecture from the HS-AFM images was challenging due to the low spatial resolution, which makes it difficult for the human eye to recognize branching points and terminals in the actin network. Additionally, the presence of AFM-specific noise, caused by the high scanning rate of the AFM needle, further imitates this method. The images were distorted with sine noise perpendicular to the scanning direction. The third difficulty is imaging of the three-dimensional (3D) F-actin network structure in a 2D representation. Individual F-actins located at different depths seemingly intersect in HS-AFM images. This makes discriminating between overlapping and branching at each cross point difficult.

Recently, a hybrid AMF imaging system was developed that can simultaneously perform HS-AFM and fluorescence imaging^14^. The system was used to successfully clarify the dynamic relationship between cellular architecture imaged using HS-AFM and molecular fluorescence signals, during endocytosis^4^. Similarly, HS-AFM imaging of actin networks can be combined with imaging using Förster (or fluorescence) resonance energy transfer (FRET)-based biosensors. For example, the spatiotemporal activity of signaling molecules such as Rac1 and Cdc42, which are Rho GTPases responsible for cell motility, can be live-imaged using FRET-based biosensors^16^. Such hybrid AMF imaging with FRET-based biosensors should improve our understanding of how intracellular signaling controls cellular morphology via the reorganization of the actin network. A methodology that extracts the topology of the F-actin network from HS-AFM images is therefore required.

In this study, we developed a new machine learning method called cytoskeletal location of various filamentous entity (cyto-LOVE) to quantitatively recognize F-actin networks in noisy, low-resolution HS-AFM images to an individual filament level. The new method can estimate the location and orientation of F-actin from noisy, low-resolution observations and track each F-actin to extract the topology of the F-actin network. By applying our method to HS-AFM images of lamellipodia and cortex F-actin, the actin network was successfully modeled from low-resolution images and morphological network attributes, such as orientation and persistence length distribution, are determined. Using cyto-love, we discovered that F-actins in the lamellipodia predominantly orient at ±35° relative to the membrane, aligning with the angle of Apr2/3-induced branching. Additionally, we found that F-actins in the cell cortex exhibit four specific orientations, challenging the widely held belief of their random orientation.

## Results

### HS-AFM imaging

HS-AFM can be used to live-image F-actin dynamics beneath the cellular membrane^15^, using a needle that is targeted to living cells to scan across the surface area of interest at a high-speed (e.g., 0.1 frames/s; **Fig. 1A**). A needle was pushed into the cell surface using a weak loading force to successfully visualize the dynamics of the F-actin in the lamellipodia at the leading edge and in the cell cortex, which is distant from the leading edge in motile cells (**Fig. 1B and C**). Although this is the first report in which F-actin dynamics have been imaged in vivo, it was difficult to recognize individual F-actins using the spatial resolution of the HS-AFM, which is determined by multiple factors, including needle width, scanning rate, and flexibility of the cellular membrane. Furthermore, the AFM image inevitably included scanning noise caused by the line scan of the needle (**Fig. 1B and C**).

**Figure 1:**
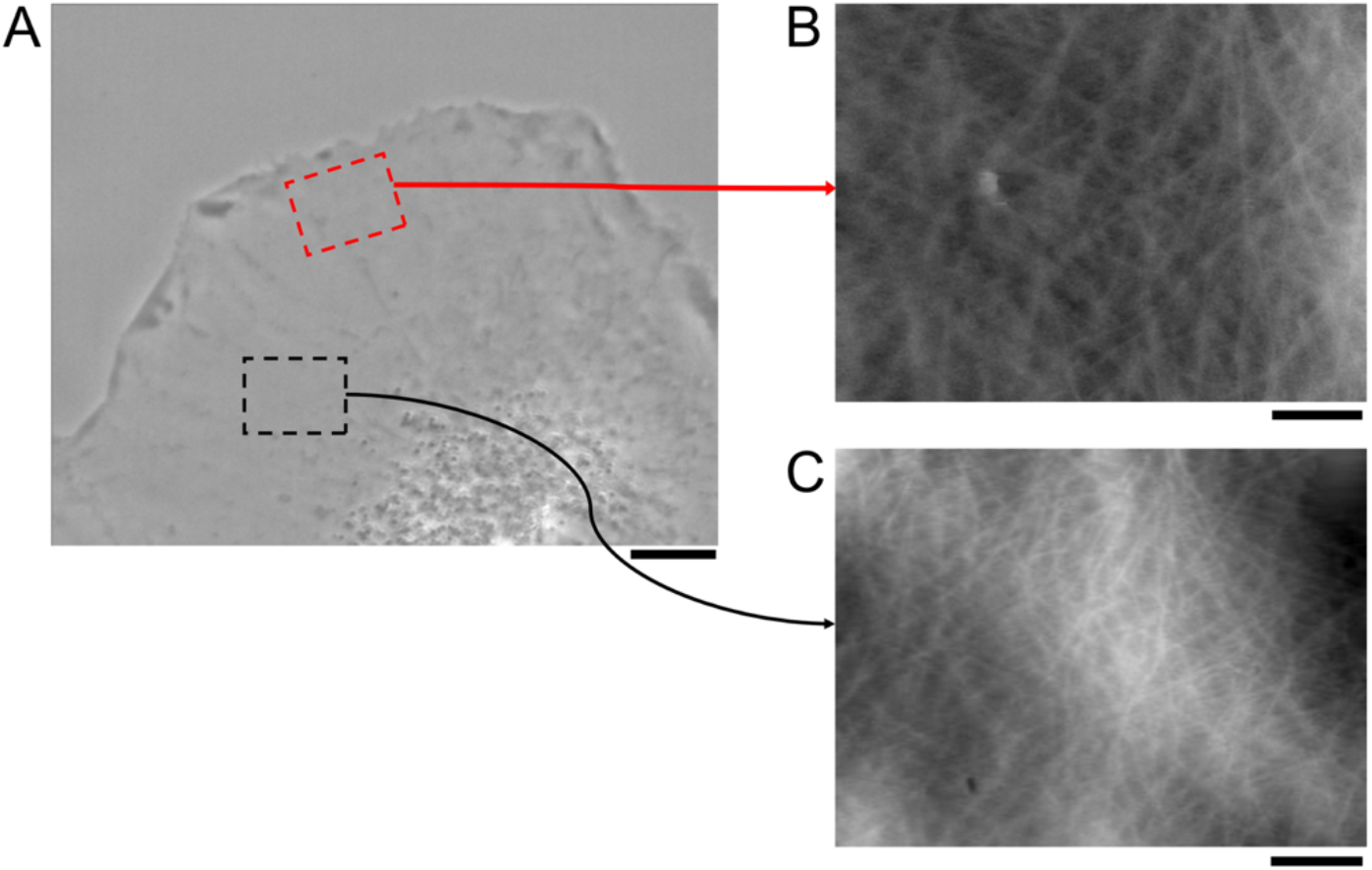
HS-AFM imaging of F-actin dynamics in living cells. **(A)** Image showing the areas of interest for the HS-AFM imaging system for live cells. Red and black dashed-line boxes represent the areas in which the F-actins in the lamellipodia and cortex were imaged, respectively. Scale bar: 3 µm. **(B)** HS-AFM images of F-actins in the cell surface area of the lamellipodia and **(C)** cell cortex. Scale bar: 1 µm.

### Data processing flow for cyto-LOVE

To quantitatively extract the topology of the F-actin network from AFM images, a new machine learning-based image analysis method was developed to determine the cytoskeletal locations of various filamentous entities (cyto-LOVE). The flow of the developed cyto-LOVE process is depicted in **figures 2A and C**.

**Figure 2:**
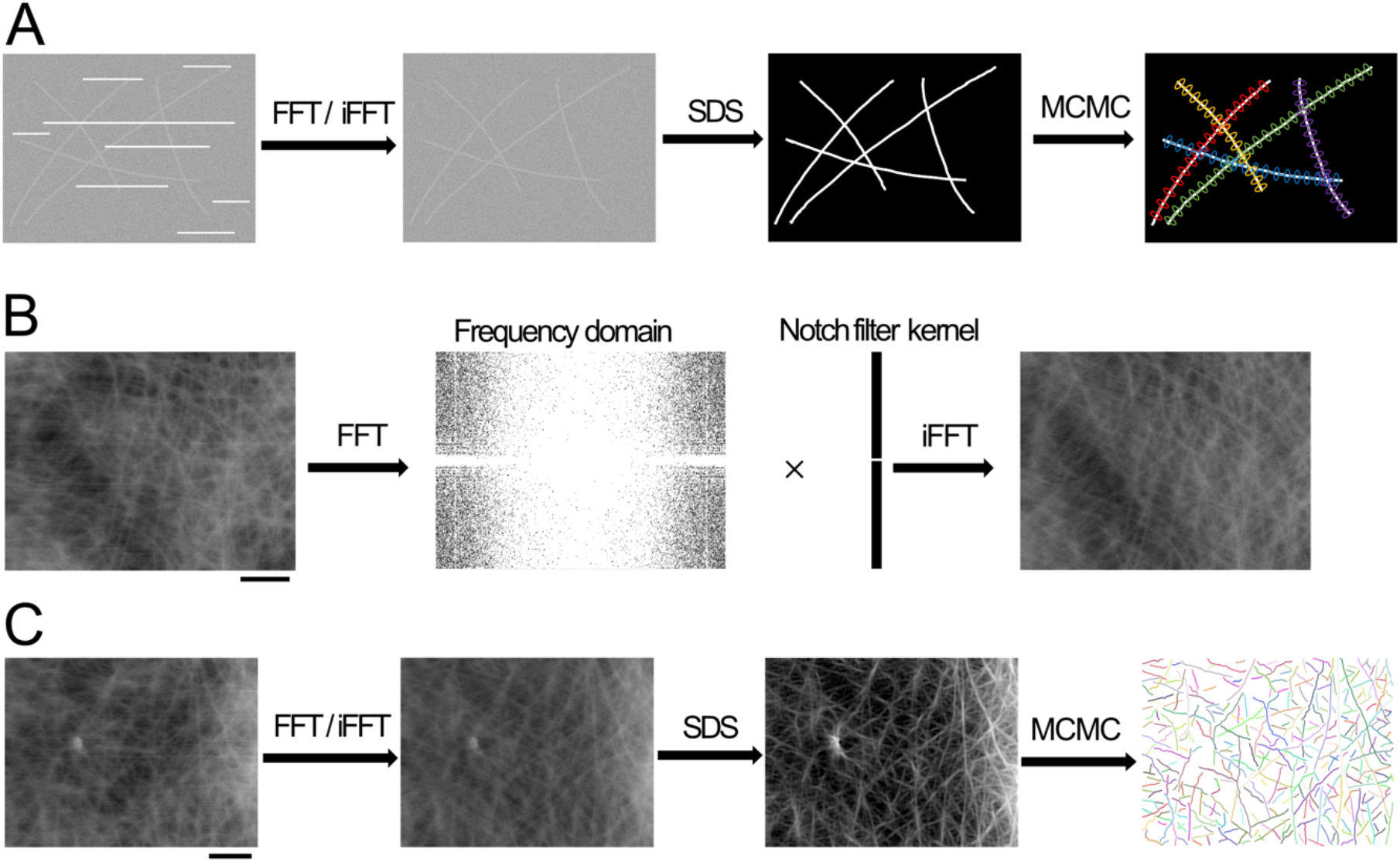
Schematic illustration of the cyto-LOVE processing flow. Flow of data processing in cyto-LOVE. FFT/iFFT processing removes scanning noise. SDS clears filamentous structures of the F-actins by estimating the presence and orientation of individual F-actins. MCMC extracts the F-actin structure as a connected particle network. The Notch filtering method is used to remove scanning noise. The HS-AFM image was transferred to the frequency domain using fast Fourier transform (FFT). The notch filter kernel was applied to remove the frequency component of the scanning noise. The denoised image is reconstructed using an inverse Fourier transform (iFFT). Scale bar: 1 µm. **(C)** Demonstration of the cyto-LOVE process using a real AFM image. Scale bar: 1 µm.

Data processing in cyto-LOVE consists of preprocessing and two essential steps. During preprocessing, scanning noise was removed by notch filtering using a fast Fourier transform (FFT; **Fig. 2B**; **Fig. 2A** after FFT/iFFT). The first step after preprocessing was the Bayesian estimation of the location and orientation of individual F-actins based on the steerable deconvolution smoothing (SDS) algorithm^20^. Noisy AFM images were clarified, indicating that the filamentous structures were highly enhanced (**Fig. 2A** after SDS). The second step was recognition of the F-actin architecture as a connected-node network by tracking the individual F-actins using the Markov chain Monte Carlo (MCMC) method ^17^ (**Fig. 2A** after MCMC).

### Step 1: Image clarification using MAP estimations of the cytoskeletal location

The first step in the cyto-LOVE process is to estimate the location and orientation of individual F-actins using MAP estimations. To represent the F-actin orientation, we have introduced a function, *Z*^∗^(***r, n***), which describes the probability of F-actin orientating at specific angles at pixel coordinate ***r*** ∈ *G* ⊂ ℝ^2^, where *S*_1_ ∋ *n* = (sin*θ*, cos*θ*)^***T***^ (*S*_1_ is the 1-sphere) denotes a unit vector with angle *θ* (−*π* ≤ *θ* ≤ *π*). This function is known as the angle orientation distribution (AOD) function^18,19^. An example of the AOD function on an image with two filaments at different angles is shown in **figure 3A**. At point (a), the AOD function peaked at 45°, as the filament running through point (a) was oriented at 45°. At point (b), the AOD function had double peaks at 45° and 90°, as point (b) is located at a cross point between two filaments oriented at 45° and 90°. However, at point (c), which is outside the filament, the AOD function had no peak and remained low.

**Figure 3:**
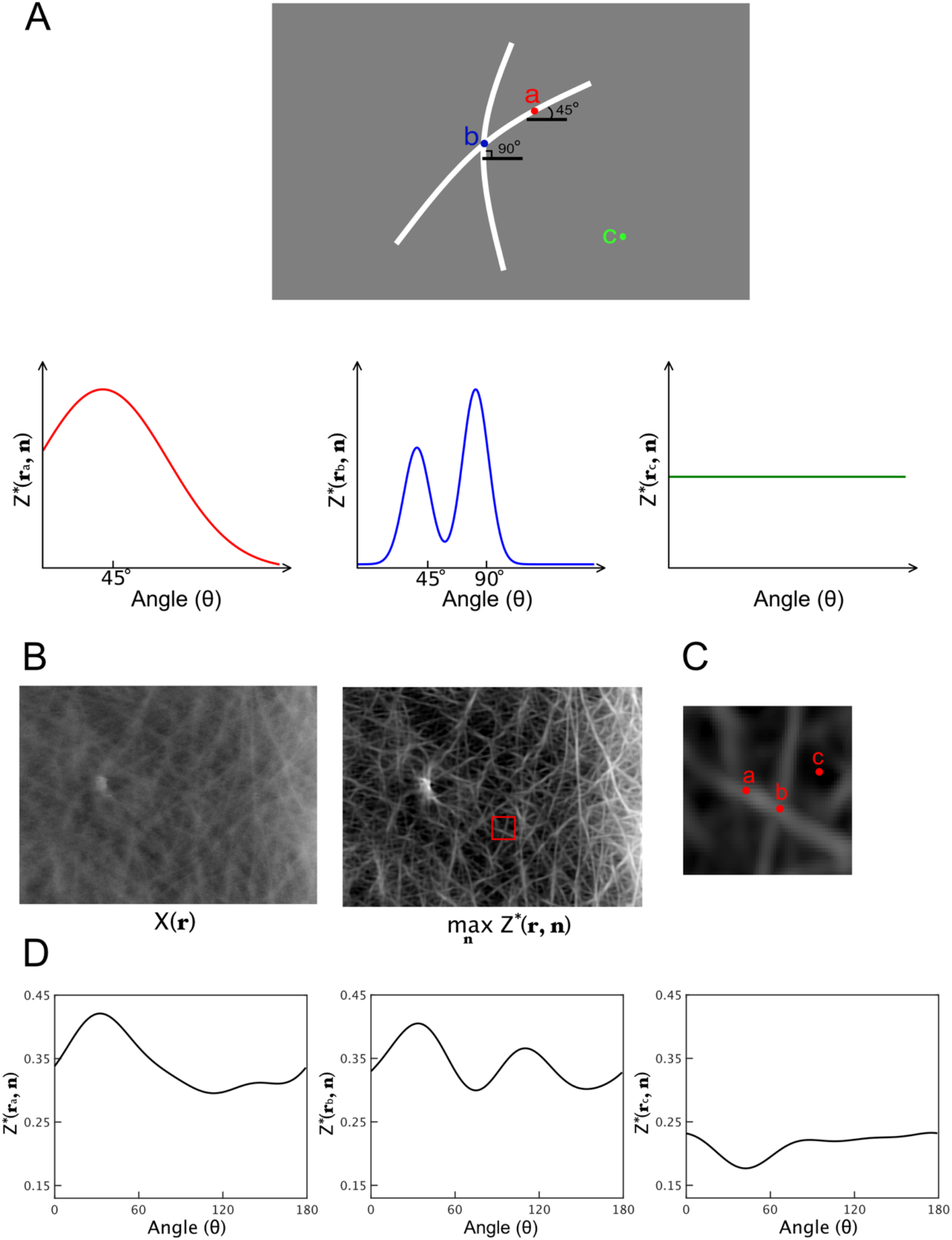
Angle orientation distribution (AOD) function for describing orientation of F-actin. **(A)** Representation of F-actin orientation when using the AOD function. The image shows an example of two crossed F-actins. The AOD functions are depicted at positions (a–c) in the image. **(B)** Clarified image derived from the AOD function estimated from an AFM image. This image was visualized using the maximum values of the AOD function with respect to *n* at each pixel. Scale bar: 1 µm. **(C)** Magnified images of the red rectangular ROI in image (B). **(D)** The estimated AOD functions are depicted at positions (a, b) in image (C). At point (a), the AOD function has a single peak that indicates a filament without crossing. At point (b), the AOD function has double peaks that indicates filaments with crossing. At point (c), the AOD function has no peak and remained low because point (c) is outside the filament.

A previously published method was used to estimate *Z*^∗^(***r, n***)^20^. According to Bayes’ theorem, the following relationship holds:

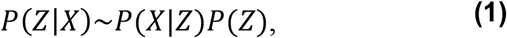

where *Z* = { *Z*(***r***_*i*_, ***n***_*i*_)}_*i*=1,2,…_ and *i* is the pixel index. *P*(*Z*|*X*) represents a posterior distribution of *Z* given image *X, P*(*X*| *Z*) represents a likelihood, and this describes a probabilistic process by which the AFM images are generated when using an unknown orientation for the F-actins; and *P*(*Z*) represents a prior distribution of *Z*, which regularizes the continuous fiber structure of F-actins (see Methods for details). This was followed by the estimation of *Z* by maximizing *P*(*Z*|*X*) as follows:

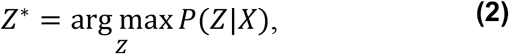

where *Z*^∗^ = {*Z*^∗^(***r***_*i*_, ***n***_*i*_)}_*i*=1,2,…_ and *Z*^∗^(***r***_*i*_, ***n***_*i*_) indicate the most likely AOD functions. The filamentous structures of the F-actins were then reconstructed (**Fig. 3B**) as follows:

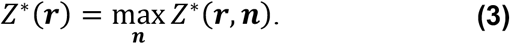

The maximization of *Z*^∗^(***r, n***) with respect to ***n*** (Eq.3) enhanced the F-actin signal, as shown in **figure 3B**. To validate the estimation of the AOD function, we confirmed that it had a single peak corresponding to the angle of the filament at the non-crossing point and double peaks corresponding to the two filaments oriented at different angles (**Figs. 3C and D**). Furthermore, it can be easily noticed that the F-actin network was visualized as if it was in three-dimension space, especially at the crossing point of F-actins (**Fig. 3B**). This was because at the maximum operation, the AOD function at the crossing points had two peaks (**Fig. 3D**), and the selection of the highest peak by the maximum operation only visualizes the F-actin with the higher intensity, which could be present near the membrane.

### Step 2: Object recognition of individual F-actins using MCMC

The second step in the cyto-LOVE method is to reconstruct the topological structures of the F-actin network using the AOD functions estimated in Step 1. As previously described^19,21^, F-actin was represented by the model *M* = {*P*, દ}, which is built upon a set of particles *P* and the set of connections દ between them (**Fig. 4A**). In this model, a particle *p* ∈ *P* is ellipse-shaped, and the orientation ***n***_*p*_ ∈ G represents the orientation of the F-actin, while position ***r***_*p*_ ∈ G represents the location of F-actin; and the edge *u* ∈ દ connecting the particles determines the topology of the F-actin network. To find a model of the F-actin network from the AFM images, a function that describes how well the model fits the F-actin network in an observed image is required. The function is then maximized by tuning the model, that is, reorganizing the F-actin network model. This function is the posterior probability of model *M* given AFM image *X, P*(*M*|*X*), which can be computed using Bayes’ theorem as

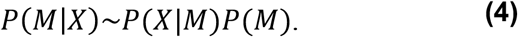

*P*(*X*|*M*) is the likelihood representing how well the model fits the observed image *X*, described by

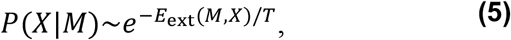

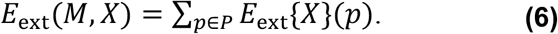

*E*_ext_(*M, X*) is called the “external energy,” where *E*_ext_{*X*}(*p*) = −*a*(*Z*^∗^{*X*}(***r***_*p*_, ***n***_*p*_) − *b*) (*a* > 0 and *b* are constants), *T* denotes the temperature parameter and *Z*^∗^{*X*} is the estimated AOD function from image *X*. Note that *E*_ext_(*M, X*) is an energy representing how the model differs from the image *X*; at the minimum of the energy landscape of *M*, the model most fits the noisy image *X* observed using HS-AMF. *P*(*M*) is the prior probability representing the connection rule of the particles in the model, which is described by:

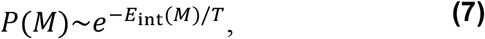

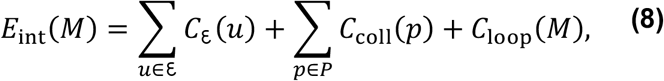

where *E*_int_(*M*) is the “internal energy,” *C*_દ_(*u*) describes the elastic energy of the F-actin,*C*_coll_(*p*) is the collision energy between the neighboring particles, and *C*_loop_ (*M*) is an energy to prohibit the loop structure of the F-actin network (see Methods in details). Therefore, the posterior probability can be written as

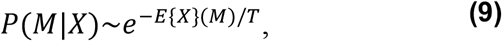

where *E*{*X*}(*M*) = *E*_ext_(*M, X*) + *E*_int_(*M*) denotes the total energy. Model *M* was estimated by maximizing *P*(*M*|*X*), which is the same problem as minimizing the total energy *E*{*X*}(*M*).

**Figure 4:**
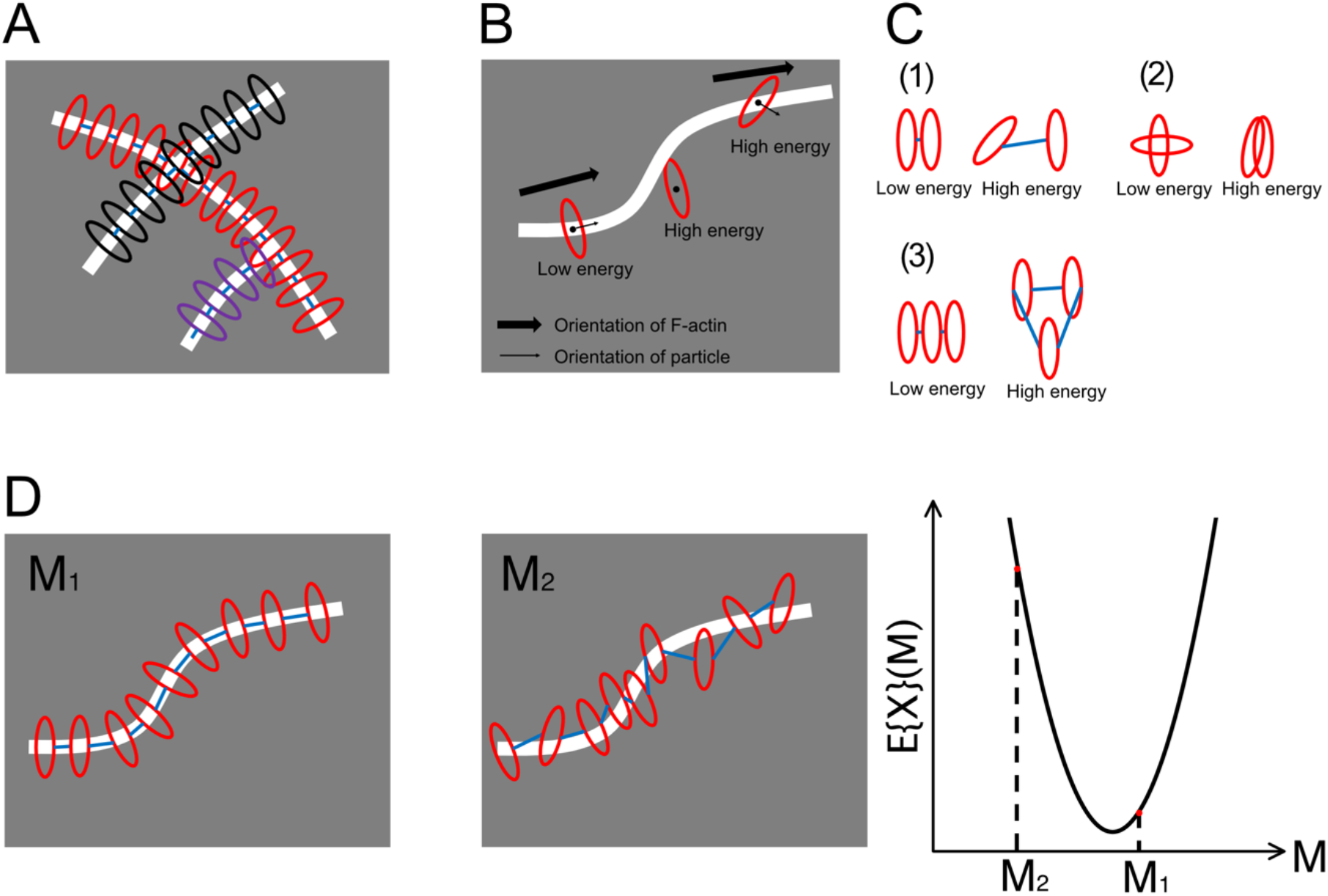
Model describing topological structure of F-actin. **(A)** Connected particle model for topological structure of F-actin. Particles are ellipse-shaped, and their semi-minor axis represents the orientation of the F-actin. **(B)** Particles in model *M* with low external energy should be placed on the central line of and orient with F-actin. **(C)** In model *M* with low internal energy, 1) two connected particles should orient at the same angle and be placed in a short distance; 2) particles can collide nearly orthogonally to represent the crossing of the two F-actins but should not collide parallelly; and 3) no closed loop appears. **(D)** Example of the total energy *E*{*X*}(*M*) of two models for a single F-actin. *E*{*X*}(*M*_2_) is lower than *E*{*X*}(*M*_1_) because *M*_”_ fits the F-actin structure in the observed image better.

The original model^19,21^ did not fully represent the F-actin network in the HS-AFM image, as its function was to track axonal fibers, whose characteristics differ from F-actins. Axon fibers are wiggly wired in the brain tissue, whereas F-actin exists in smooth and continuous segments with a high persistence length. In the original formulation, *C*_દ_(*u*) was described by a single energy term simultaneously that included the repulsive and bending effects of particles. However, the effect of bending was weak compared with the repulsive effect and cannot be controlled independently of the repulsive effect. Therefore, cyto-LOVE introduces a new internal energy *C*_દ_(*u*) that strengthen the bending of the connected particle to represent the smooth and continuous segments of the F-actin network (see Methods)

The estimated model must satisfy three characteristics: 1. elliptical particles should be placed on the central F-actin line; 2. particles should be oriented along the F-actin; and 3. the two adjacent particles should be placed at almost equal distances. The external energy, *E*_ext_(*M, X*), is minimized when a model satisfies the first and second characteristics, indicating that particles are accurately aligned with the F-actin’s central line and orient along it (**Fig. 4B**). This alignment suggests a strong correspondence between the model and the observed data. The internal energy, *E*_int_(*M*), is reduced when the model satisfies the third characteristic, specifically when two connected particles maintain the same orientation at a close distance ([1] in **Fig. 4C**). However, the internal energy increases significantly when particles either collide while parallel ([2] in **Fig. 4C**) or form a closed loop ([3] in **Fig. 4C**).

Notably, the *E*_int_(*M*) was designed such that it has a low value when two particles collide in a nearly orthogonal manner, representing the crossing of two F-actins ([2] in **Fig. 4C**). Consequently, the more model *M* meets the three identified characteristics, the lower the total energy value *E*{*X*}(*M*) (**Fig. 4D**).

We optimized model *M* by minimizing *E*{*X*}(*M*) using the RJMCMC method^17^. In each iteration of the RJMCMC method, an alternative model, *M*_new_, is sequentially generated from the current model in a random manner and then probabilistically accepted or rejected (see Methods). Alternate models are then generated as follows; particle birth and death (adding or deleting particles from the current model; **Figs. 5A and B**), moving and rotating of particles (changing the position or orientation of particles; **Fig. 5C**), connection/reconnection between particles (adding or altering connections; **Fig. 5D**), and track-birth and track-death (like the polymerization and depolymerization of actin, adding or removing particles from the end of a modeled F-actin; **Figs. 5E and F**).

**Figure 5:**
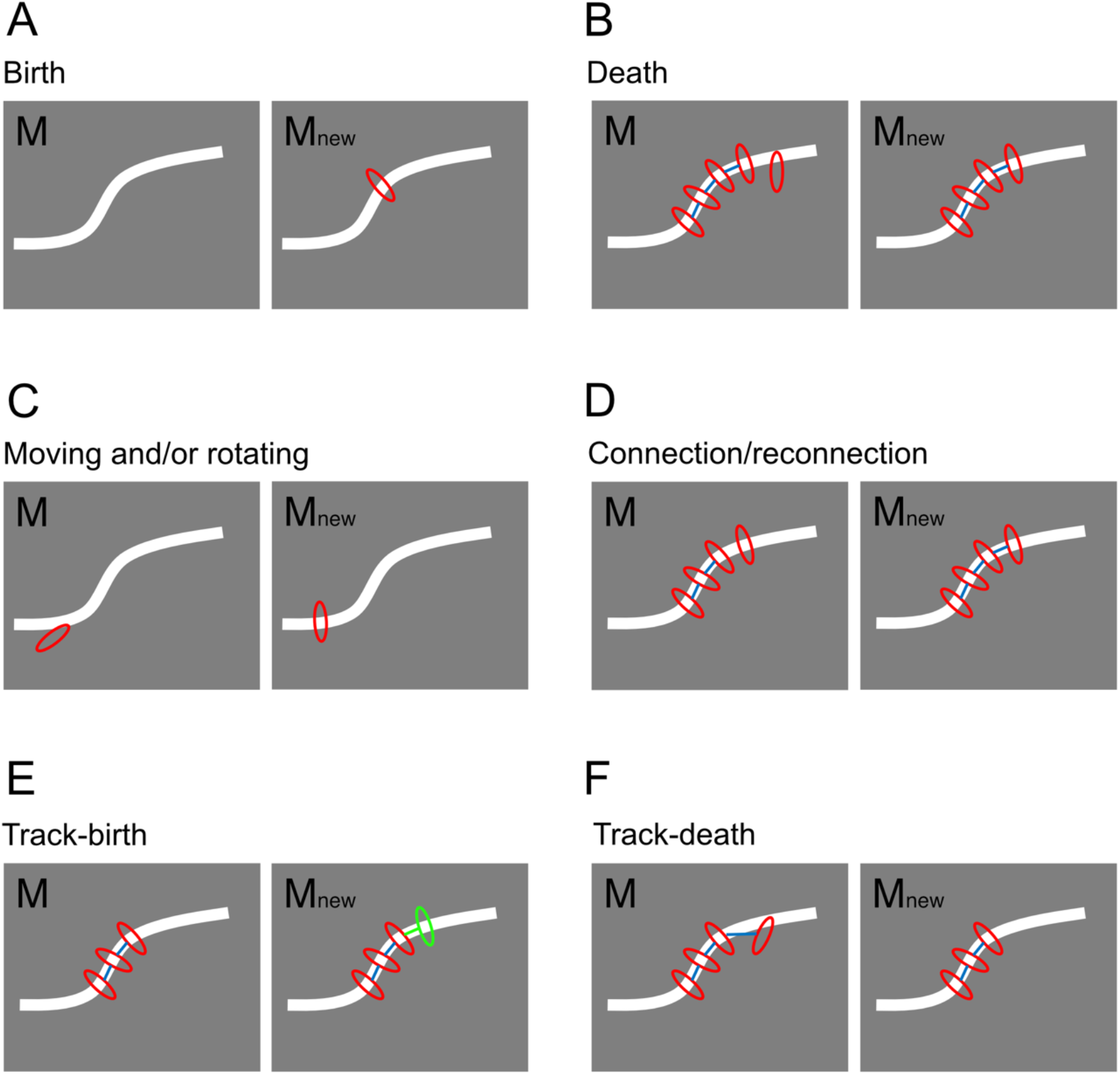
Efficient sampling of the MCMC method to optimize the connected particle model. **(A)** Particle birth; a new particle is randomly added to the current model. **(B)** Particle death; a particle is randomly selected to be removed from the current model. **(C)** Moving and rotating of particles; the positions or orientations of the particles are altered. **(D)** Connection/reconnection between particles; a new connection is created between two adjacent particles or an existing connection is reconnected to another adjacent particle. **(E)** Track-birth of particles; F-actin is randomly selected, and a new particle is added to the end, similar to polymerization. **(F)** Track-death of particles; F-actin is randomly selected and a particle at the end is removed, similar to depolymerization.

To test the performance of the MCMC method, it was used to process an artificial image created using two independent crossing F-actins and Gaussian noise (**Fig. 6A**). Model estimations were then visualized during the MCMC iterations (**Fig. 6B**). Here, we observed the elementary model alterations in the MCMC. First, seed particles are generated and randomly placed on the F-actin by “birth” (plotted in red color; at 100 iterations in **Fig. 6B**). Based on those particles, “track-birth” generates particles (plotted in green color) to elongate the connections like the polymerization of F-actin (at 500 iterations in **Fig. 6B**). On the same F-actin, two separated connections appeared (at 1,000 iterations in **Fig. 6B**), as the seed particles were randomly generated by “birth.” The two separated connections were elongated by “track-birth,” then met head-on and linked by “connection/reconnections” (at 10,000 iterations in **Fig. 6B**). At this moment, the MCMC recognized the crossing of two independent F-actins, not as an x-shaped object (at 10,000 iterations in **Fig. 6B**; **Fig. 6C**), as the collision of differently oriented particles was allowed at the crossing point (**Fig. 6D**; see Methods). Finally, the F-actin networks were fully represented by the connections between particles (at 50,000 iterations in **Fig. 6B**). The total energy of the model was confirmed to decrease during the MCMC iterations (**Fig. 6E**).

**Figure 6:**
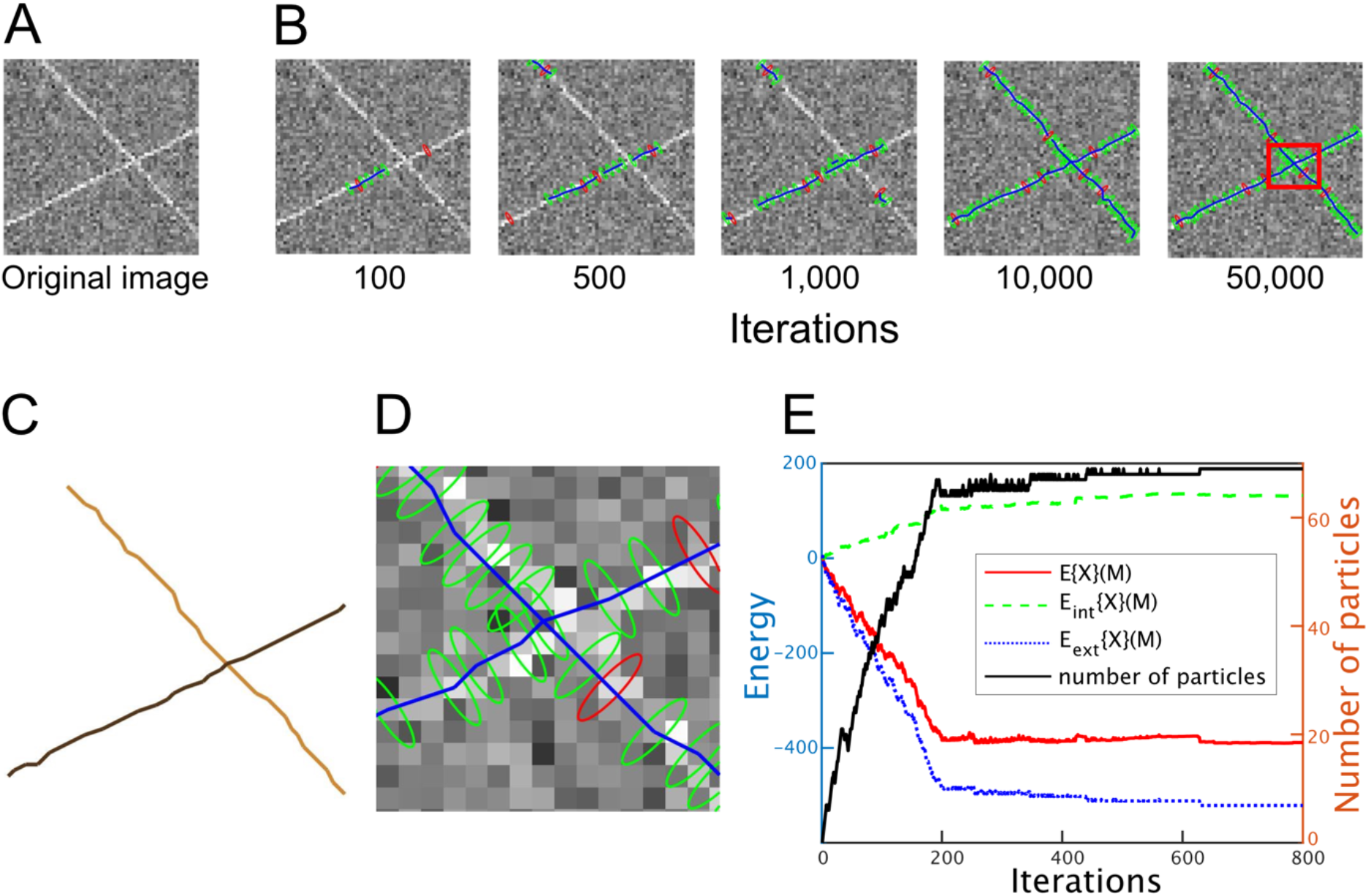
Demonstration of the developed method using an artificial F-actin image. **(A)** Artificial image containing two crossing F-actins. Noise was added to the image, which mimics the actual AFM images. **(B)** Visualization of the MCMC method. The F-actin network is represented by connected-ellipse-shaped particles, and the green and red particles are generated by the track-birth and birth samplings, respectively. **(C)** Estimated model of the F-actin network. **(D)** Magnifications of the red ROI at 50,000 iterations in (B). Note the colliding particles, each of which are assigned to different F-actins. (E) Change in the total energy during the MCMC iterations. Note that the intrinsic energy *E*_int_{*X*}(*M*) consisting of the F-actin elastic energy *C*_દ_(*u*) and the collision energy *C*_coll_(*p*) has a positive value as the number of particles increases, while the external energy *E*_ext_{*X*}(*p*) always has a negative value.

### F-actin network characteristics in the lamellipodia and cell cortex

The cyto-LOVE method was used to process an HS-AFM image of COS-7 cells in culture (**Fig. 7**). To observe the subcellular F-actin structures, HS-AFM was used to image the lamellipodia and the cell cortex distant from the lamellipodia.

**Figure 7:**
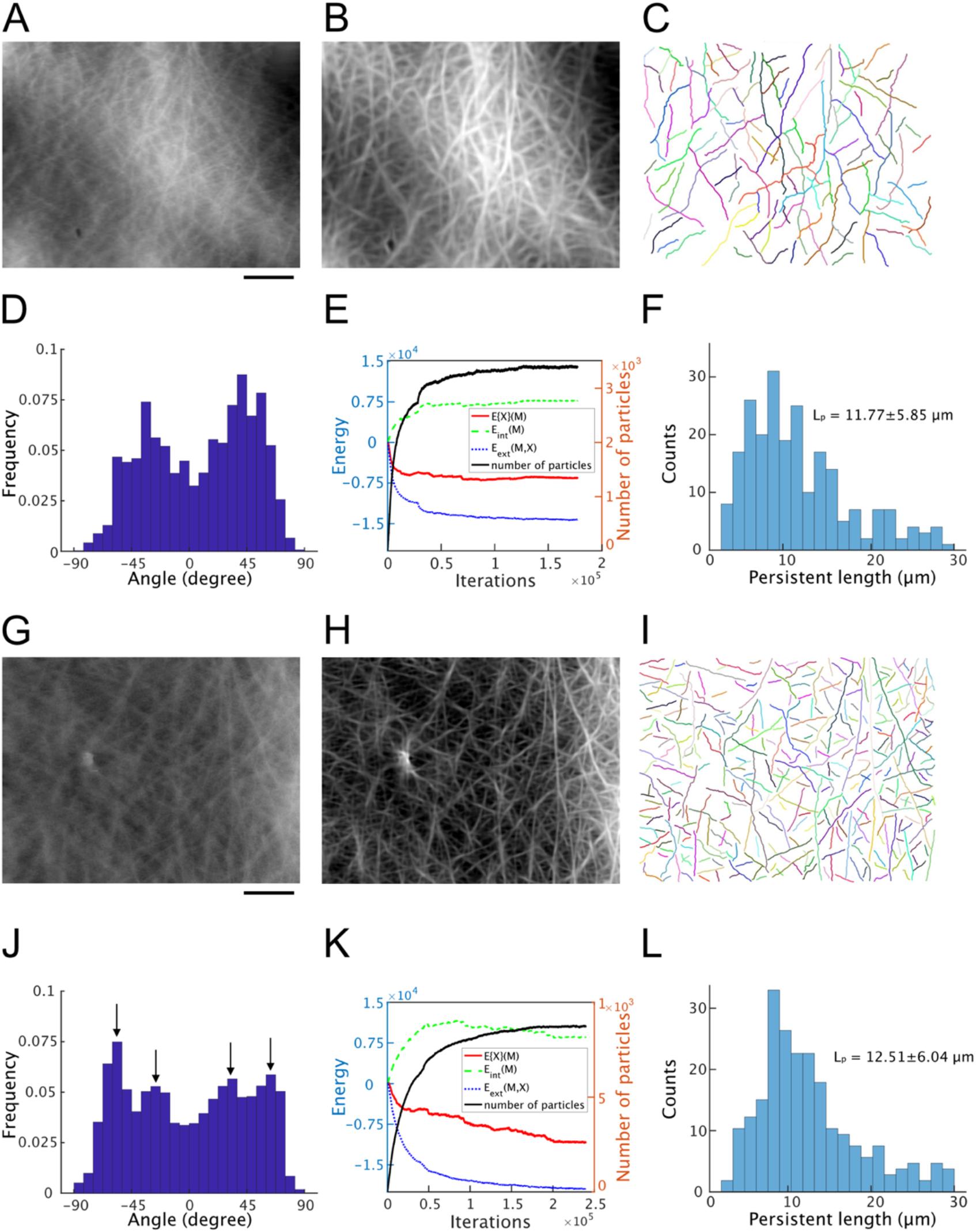
Different orientations of estimated F-actins in the lamellipodia and cell cortex. **(A, G)** AFM images of the lamellipodia and cell cortex. The image contains a large amount of scanning noise along the scanning direction of the AFM needle. Scale bar: 1 µm. **(B, H)** AFM images after processing using the FFT/iFFT and SDS algorithm. The scanning noise in (A, G) was removed using FFT/iFFT. The image was visualized using the maxima of the estimated AOD function with respect to angle at each pixel. **(C, I)** Estimated F-actin structures. Individual F-actins are depicted in different colors. **(D, J)** Distribution of the estimated F-actin angle structures in the lamellipodia and cell cortex. The individual F-actin angles were calculated using the average angle of the connecting particles. (**E, K**) Changes in the total energy during the MCMC iterations in the lamellipodia (E) and cell cortex (K). (**F, L**) Persistence length distribution of lamellipodia (F) and cell cortex (L) network. The persistence length *L*_p_ is measured to be 11.77 ± 5.85µm and 12.51 ± 6.04µm respectively.

The AFM images of the lamellipodia were examined first. For the cyto-LOVE data processing, scanning noise was removed from the raw images using FFT/iFFT methods (**Fig. 7A**); the images were then clarified using the SDS algorithm (**Fig. 7B**), and the F-actin structures estimated using the MCMC algorithm (**Fig. 7C**). The angles of individual F-actin were quantified in the estimated F-actin structure. It was then confirmed that the angle distribution had two peaks in the lamellipodia, with an interpeak interval of 70°, which was consistent with the angle of branched F-actin by the Arp2/3 complex ^6^ (**Fig. 7D**). These results clearly indicate that cyto-LOVE can accurately estimate the F-actin structure. The total energy of the model was confirmed to decrease during the MCMC iterations (**Fig. 7E**). The persistence length *L*_p_ is measured to be 11.77 ± 5.85µm in the estimated F-actin structure and the distribution was shown in **figure 7F**. The persistence length of F-actin was reported to be nearly 16.7µm with low deviation when its length ranges between 5 and 15µm^22^.

The AFM images of the cell cortex were then examined using the cyto-LOVE method (**Figs. 7G–L**). The F-actins in the cell cortex were non-randomly oriented, which is in contrast to that in previous studies that reported that they were uniformly oriented^23,24^. In contrast to the two peaks in the lamellipodia, the angle distribution had four peaks at approximately −60°, −30°, 30°, and 60°, respectively, in the cell cortex (arrows in **Fig. 7J**). Two orthogonal pairs (−60° and 30°, and −30° and 60°) were identified and indicated the presence of an actin-associated protein, filamin A (FLNa), which orthogonally links two F-actins in the cell cortex^25^. Furthermore, this strongly suggests a novel mechanism for F-actin organization that involves FLNa, which exists latently in the cell cortex.

## Discussion

In this study, a machine learning method (cyto-LOVE) was developed to quantitatively recognize F-actin networks in HS-AFM images. The method sequentially conducts a two-step computation: the first step extracts the angle orientation of F-actin as the AOD function, and the second step recognizes the individual F-actin represented as the connected particle model, which is biologically reasonable and fits well with the observed HS-AFM images using MCMC. By analyzing the HS-AFM images of COS-7 cells, we demonstrated that cyto-LOVE can accurately reconstruct the Arp2/3-indued branched F-actin network in lamellipodia. Moreover, we identified novel characteristic of F-actins in the cell cortex, specifically that F-actin dominantly orients at four specific angles, although it is widely believed that F-actins are uniformly oriented in this region. This discovery suggests the existence of an unknown mechanism for F-actin organization. The developed cyto-LOVE method will thus be a fundamental tool to help improve our understanding of the structural dynamics of the F-actin network.

The approach described in this study has three defining characteristics when compared with previous models. First, cyto-LOVE is robust in its ability to recognize individual F-actins in noisy AFM images with low spatial resolution. This robust ability was accomplished by estimating the AOD function, in which prior knowledge regularized the shape of F-actin as a continuous fiber structure. In a low-resolution image, noise can generate random patterns rather than a specific continuous fiber structure. Thus, prior knowledge of the F-actin structures enables a more robust estimation of AOD function by suppressing the noise effects. Second, cyto-LOVE adopts the biological kinetics of F-actin to optimize the connected particle model. In the MCMC, track-birth and track-death proposals were used for the sampling models of the F-actin network. The track-birth proposal mimics the polymerization of F-actin by adding a new particle at the terminus using a single particle as an actin monomer. The track-death method mimics depolymerization by removing the terminal particles. Thus, these proposals efficiently sampled biologically realistic models of the F-actin network in MCMC. Third, cyto-LOVE adopts the physical properties of F-actin, which is a stiff and almost straight filament that has a high level of persistence. In MCMC, we incorporated a novel bending elastic energy to realize a realistically smooth and continuous F-actin structure.

However, two problems still need to be addressed. The first is time-lapse tracking of individual F-actins. It is difficult to predict what a certain F-actin will become in the next frame during time-lapse tracking, as their length, position, and shape change between neighboring frames and they may even disappear. The second is the recognition of branching points, as owing to the low resolution of HS-AFM images, they are not recognized by the human eye. Future research will thus be required to improve the resolution of the HS-AFM images in the experiment and develop a model that includes branching points in theory.

The tracking of individual F-actins using cyto-LOVE revealed that they predominantly orient at ±35° respective to the membrane in the lamellipodia. This is consistent with the results of a previous study using EM^6,7^. In addition, the Arp2/3 complex induces a new branched F-actin at 70°, and the branched F-actin networks are specifically oriented toward the membrane at ±35°^6^. This has validated the accuracy of cyto-LOVE as a method to quantify F-actin orientation.

In the cell cortex, F-actin was thought to be randomly oriented^23,24^, but the results of this study showed that it is predominantly oriented to the membrane at four specific angles at ±30° and ±60°. Pairs of −60° and 30° and −30° and 60° correspond to the orthogonal crossing regulated by FLNa^25^. While how F-actins are organized and oriented to the four peaks is unknown, a mechanism for this phenomenon has been suggested. As a premise, it was assumed that F-actins orienting at 0° are present in the lamellipodia of non-migrating cells^26^, whereas those orienting at ±35° are regulated by Arp2/3. If F-actins orienting at 0° and −35° are linked with FLNa (green triangle in **Fig. 8**), they can reorient at −62.5° and 27.5°, which is similar to that at −60° and 30° (**Fig. 8A**). However, if F-actins orienting at 0° and 35° are linked with FLNa, they can reorient at −27.5° and 62.5°, which is similar to that at −30° and 60° (**Fig. 8B**). Further experiments, such as the knockdown of FLNa, are required to verify the proposed mechanism.

**Figure 8:**
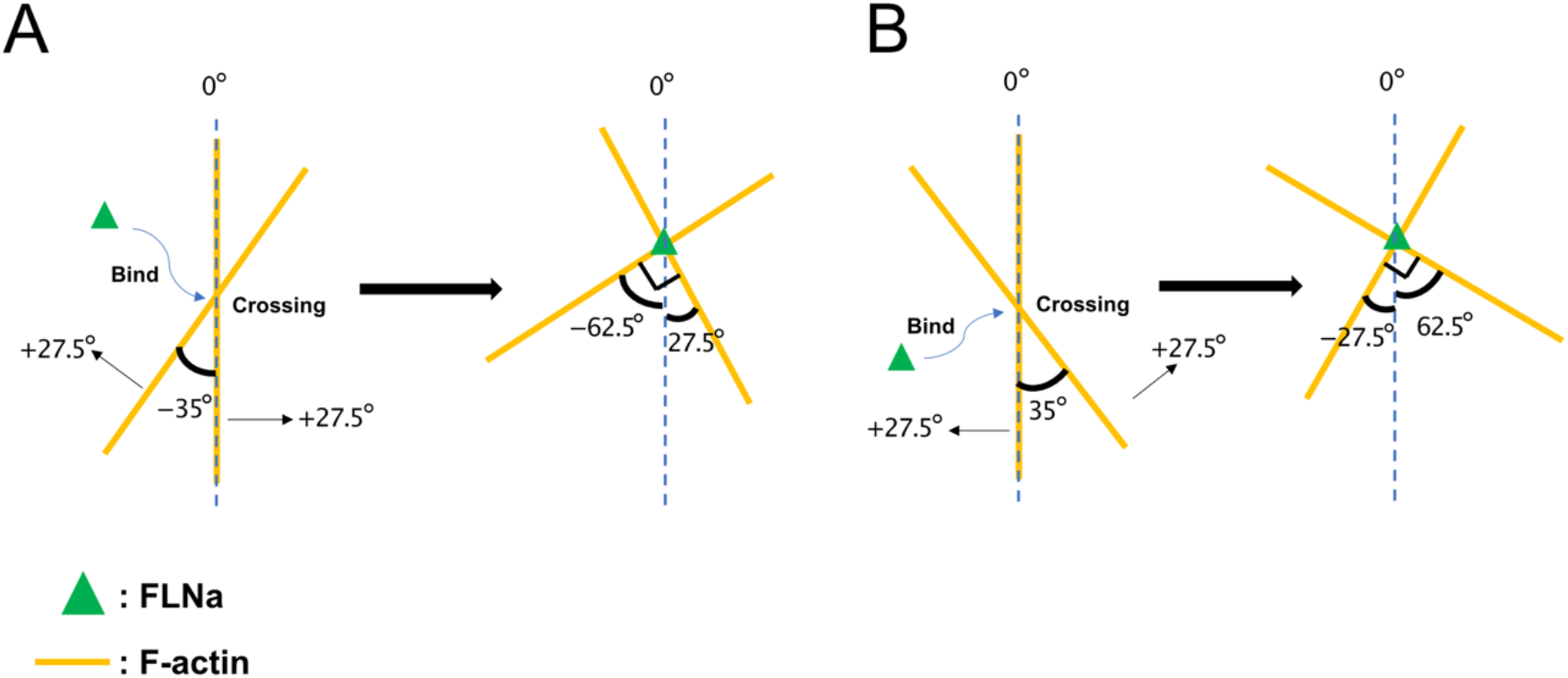
Possible F-actin organization mechanism. **(A)** Two crossing F-actins (yellow lines) with angles of 0° and −35°. Two F-actins will reorient at −62.5° and 27.5° when linked with FLNa (green triangle), which expands their angles by 27.5°, respectively. **(B)** Two crossing F-actins with angles of 0° and 35°. Two F-actins will reorient at −27.5° and 62.5° when linked with FLNa, which expands their angles by 27.5°, respectively.

The recognition and tracking of F-actins has been computationally addressed based on two methods: the Canny method following a steerable filter ^27^ and the Radon transform-based method^12^. However, these methods have two major limitations. First, they are based on direct computation of image intensities; therefore, the results are susceptible to inevitable noise in the images. Second, these methods are not applicable to low-resolution AFM images because of miss and wrong misrecognition due to a lack of prior knowledge of the F-actin structures. However, our method overcomes the shortcomings of the two methods by using MAP estimation of the AOD function from the AFM images with prior knowledge of the F-actin structure (see Eq.13 and 14 in the Methods section).

Recently, two robust methods, EMILOVE ^21^ and PAT^19^, were developed to track axon fibers. Similar to the method presented in this study, these methods use the MCMC method to reconstruct a connected particle model that represents the fiber structure. The reconstruction process of the connected particle model in these two methods was optimized for axon fiber tracking but was not suitable for F-actin tracking. This is because axon fibers are wiggly wired in brain tissues, whereas F-actins exist as smooth and continuous segments with a high persistence length. Thus, these two methods are not suitable for the analysis of F-actin images obtained using HS-AFM. The method developed in this study overcomes the shortcomings of introducing new elastic energies and developing polymerization kinetics-based MCMC samples.

Recently, we succeeded in simultaneously imagining live F-actin dynamics and molecular fluorescence signals by developing a hybrid system of HS-AFM and fluorescence imaging^4,14^. Using this hybrid imaging system with a FRET-based biosensor, it is possible to improve our understanding of how signaling molecules such as Rac1 and Cdc42 control cellular morphologies via the reorganization of F-actin networks. In addition, simultaneous imaging of molecules with unknown functions could contribute to the discovery of their functions in the organization of F-actin dynamics. To this end, the newly developed cyto-LOVE method provides a fundamental tool that will help to understand such mechanisms within the cell.

## Methods

### HS-AFM imaging

COS-7 cells were grown on a poly L-lysine-coated glass slide 1 d prior to AFM imaging at 37°C with 5% CO2 in Dulbecco’s modified Eagle’s medium (DMEM), supplemented with 10% fetal bovine serum (FBS, Hyclone). AFM imaging was performed in DMEM supplemented with 10% FBS and 10 mM HEPES-NaOH (pH 7.0–7.6; Sigma Aldrich) with a tip-scan type HS-AFM unit combined with an inverted optical microscope (BIXAM™, Olympus Corp.). An electron-beam deposited sharp cantilever tip with a spring constant of 0.1 N/m (USC-F0.8-k0.1, Nanoworld) was used. The AFM tip was targeted to a specific area of the cell based on the phase-contrast image. AFM images were then acquired at a scanning rate of 0.1 frames/s.

### Estimation of AOD functions

The AOD functions were estimated from the AFM images using the Bayes theorem Eq.1. The AOD function of the AFM image *X*(*r*) was denoted by *Z*{*X*}(*r, n*), where *r* ∈ *G* ⊂ ℝ^2^ and *n* ∈ *S*_1_ (*S*_1_ is the 1-sphere) indicate the location of the pixel and F-actin orientation, respectively. The likelihood of *X* was described using a Gaussian distribution with mean *K Z* and covariance matrix ∑, as follows:

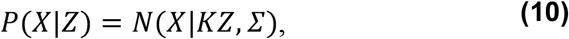

where *X* = {*x*_*i*_}_*i*=1,2,…_, *Z* = { *Z*(*r*_*i*_, *n*_*i*_)}_*i*=1,”,…_, and ∑ = *I. x*_*i*_ indicates the intensity of pixel *I i*n the AMF image. For simplicity, *X* and *Z* are used instead of *Z*{*X*}(*r, n*) and *X*(*r*) in Eq. 10. It is also of note that *K Z* in Eq. 10 represents the linear mapping from the hidden fiber structure to the AFM observation, as follows:

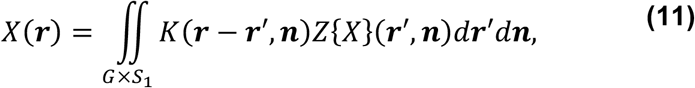

where *K:* ℝ^2^ *× S*_1_ *→ ℂ* indicates the kernel for linear mapping, and it was chosen to be imaginary and combined with both the fiber and edge detectors ^20,28^, as follows:

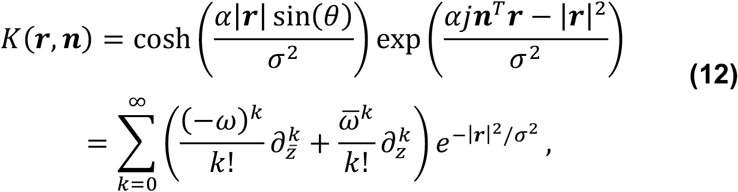

where *ω* = *e*^*jθ*^ andj is the complex number, *θ* ∈ [−*π, π*] is related to as *S*_1_ ∋ ***n*** = (cos*θ*, sin*θ*)^3^. The second line represents the Fourier series, by which the computational cost *w*as reduced, where 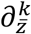 and 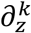 denote the *k*^*th*^ order complex derivative operator. The real part of *K* is a tubular structure detector, and the resulting shapes with various *θ* values are shown in **figure 9**. In this study, only the real part is used to estimate the AOD function. In the implementation, the summation of the Fourier series was computed until *k* = 13.

**Figure 9:**
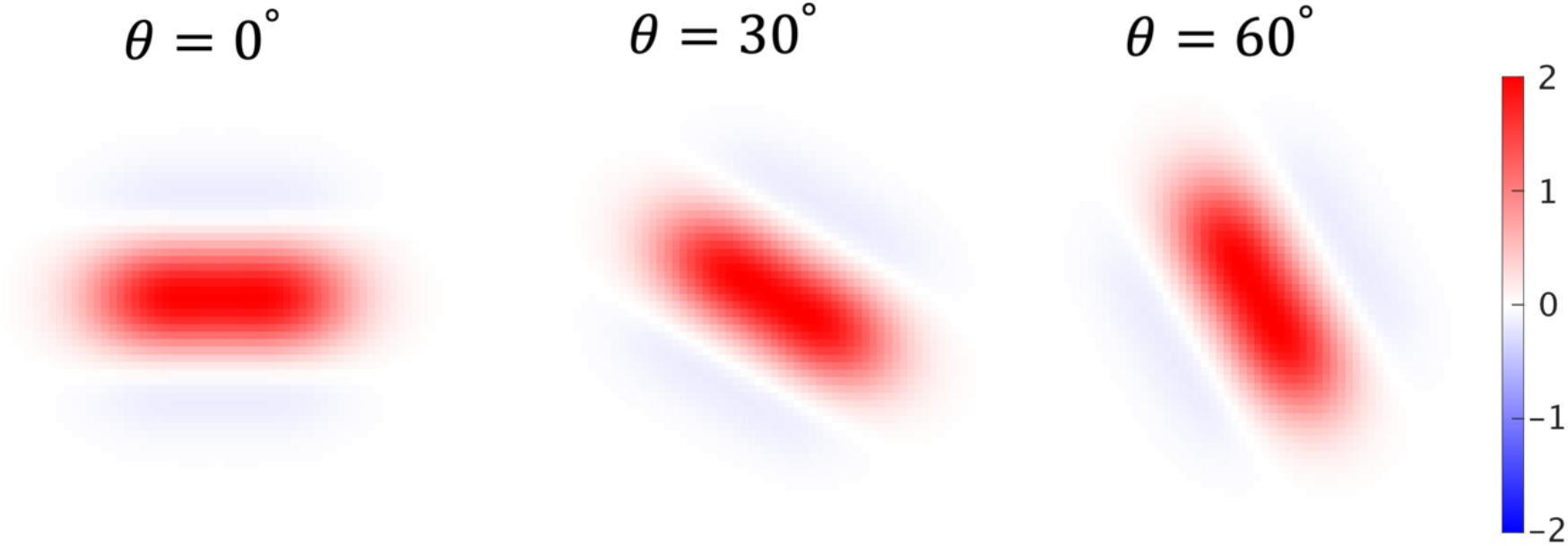
Shape of the linear mapping kernel. Shape of the real part of the kernel described in Eq. 12 with *θ* = 0°, 30° and 60°. The real part of Eq. 12 is the tubular structure detection, which can detect tubular structure based on the *θ* orientation.

A previously proposed prior distribution^18,20^ was adopted, which regularizes the continuity of F-actin as follows:

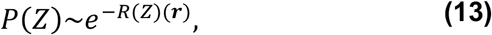

where

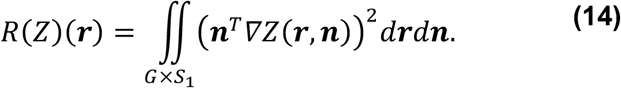

This prior constrains the F-actin, whose orientation is perpendicular to the image gradient of *Z*.

Then, *Z* was estimated using an MAP estimation. By substituting Eq. 13, 14, and 10 into Eq. 1, the maximization of the posterior of *Z* becomes the minimization of the following objective function:

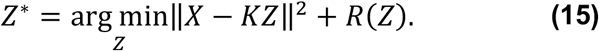

As the optimization of Eq. 15 is quadratic, the conjugate gradient descent was used to solve the problem^20^.

### Internal model energy

Here we describe *C*_દ_(*u*), *C*_coll_(*p*) and *C*_loop_(*M*) that form *E*_int_(*M*) in Eq. 8; where model *M* is represented by the two attributes of particle *p* ∈ *P*, position *r*_*p*_ ∈ *G* and orientation *n*_*p*_ ∈ *S*_1_, and where scale (*s*) and thickness (*w*) are constants 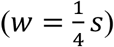 (**Fig. 10A**). In the model, particles have two connecting sites at 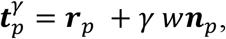 where *γ* ∈ {1, −1}, which indicates both polar sides of the particles, and that they are connected by the edge *u*. An edge *u* ∈ દ connecting two particles *p* and *p*^*’*^ is defined by 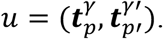 In this study, the scale and thickness of each particle in model *M* was empirically set to a fixed value of *s* = 2 and *w* = 0.5, as the individual F-actins in the HS-AMF image had almost the same thickness.

**Figure 10:**
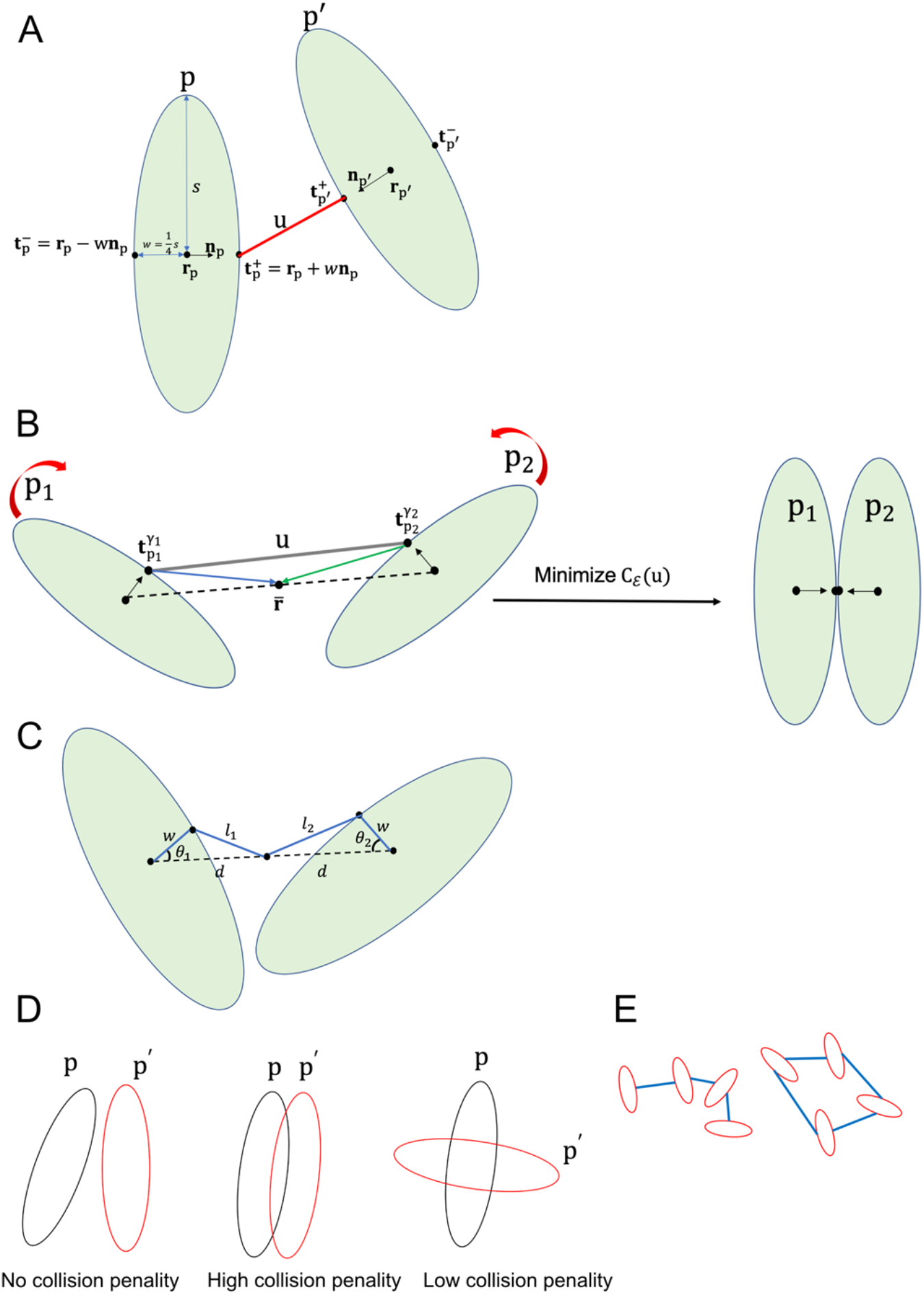
Graphical descriptions of the internal energies. **(A)** Definitions of the attributes of ellipse-shaped particles. A particle *p* is defined by its position *r*_*p*_ and orientation *n*_*p*_, where scale (*s*) and thickness (*w*) are constants shared among all the particles. The edge (*u*) is depicted by the red line and defined as follows, 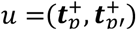, where 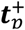 and 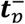 are coordinates of the particle connecting sites. **(B)** Elastic energy of the F-actin represented by Eq. 16. The pulling force is generated so as to minimize the squared sum of lengths of the blue and green lines (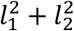), which are respectively segments from the midpoint between the central points of *p*_1_ and *p*_2_ (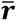) to the particles’ connecting sites (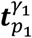 and 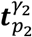). The red curve arrows represent the additional bending energy term described in Eq. 16 to strengthen the elastic penalty by penalizing the differences between the orientations of *p*_1_ and *p*_2_. **(C)** Geometric interpretation of the bending energy of F-actin represented by Eq. 16. **(D)** Collision energy between two particles. The left, middle, and right show no collision energy between the non-overlapping particles, high collision energy between the overlapping particles, and low collision energy between the orthogonally overlapping particles, respectively. **(E)** Loop energy. There is no loop energy without a closed loop (left). The loop energy is infinite with a closed loop, which means that a model with a closed loop should be discarded (right).

#### *F-actin elastic energy C*_દ_(*u*)

*C*_દ_(*u*) describes the elastic energy of the edge connecting particles (**Fig. 10B**). The energy is formulated as follows:

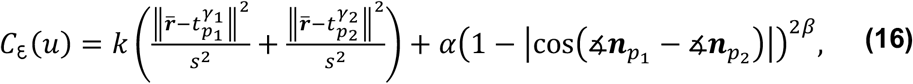

where 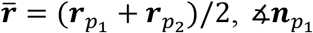 and 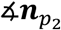 denote the angles of 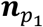 and 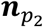, respectively, *k* and *𝛼* denote parameters weighting two elastic energies, and *𝛽* denotes the positive constant. The first term, represents the elastic energy identified in previous studies^19,21^. However, by using only this term, this energy was found to have a small effect on penalizing particles with different orientations. Consequently, bending energy was also introduced into the second term.

To illustrate the bending effect of *C*_દ_(*u*), Eq. 16 was expanded with geometric interpretation as follows:

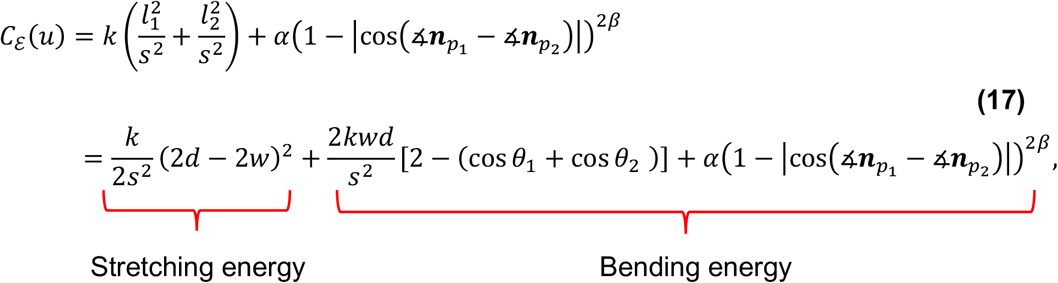

*w*here *l*_*1*_, *l*_*2*_, *θ*_1_, *θ*_2_, *d* and *w* are defined in **figure 10C**. From the first to second line, the law of cosines was used, which is as follows: 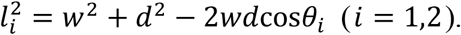 The *C*_*E*_*(u*) was then decomposed into the stretching energy in the first term and two bending energies in the second and third terms. The stretching energy penalizes the distance between the particles, and the bending energy penalizes the different angles between the connected particles. The bending energy in the second term is weak as it independently depends on the particle angles, whereas the bending energy introduced in the third term is strong as it is determined by the relative angle between two connected particles i.e.,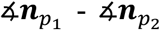. Owing to the new energy term, we can adjust the strength of the bending energy of the connected particle by setting the value of *𝛼* and *𝛽* to represent smooth and continuous segments of the F-actin network.

#### Collision energy C_coll_(p) and loop energy C_loop_(M)

*C*_coll_ (*p*) describes the collision energy between the neighboring particles^19^, the formula for which is as follows:

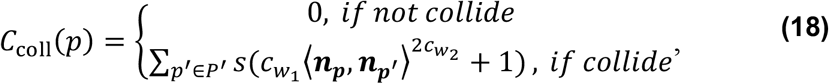

where *<*., . > denotes the inner product of two vectors, *P*^*’*^*d*enotes the set of particles that collide with *p*, and *c*_*w1*_, *c*_*w2*_ ∈ ℝ are constants. Non-overlapping particles were not penalized (**Fig. 10D** left). The inner product of this equation indicates that overlapping particles with parallel orientations are strongly penalized (**Fig. 10D** middle), whereas overlapping particles with orthogonal orientations are weakly penalized (**Fig. 10D** right). This collision energy thus allows for crossing between the two F-actins.

The loop energy *C*_loop_(*M*) prevents the loop connection of particles in the model by returning to infinity when a loop appears and 0 if there is no loop^19^, as shown in **figure 10E**:

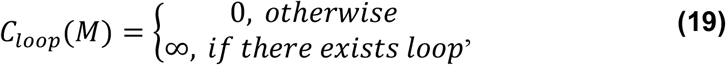

An infinite value of energy indicates that model *M* is unexpected and is discarded immediately.

### MCMC modeling method

Model *M* was optimized by maximizing *P*(*M*|*X*) using the reversible jump Markov chain Monte Carlo (RJMCMC) sampler^17^. The RJMCMC sampler is an iterative algorithm combined with simulated annealing over the temperature *T*. Specifically, we set the initial temperature at *T* = 1, and then decrease it slowly by *T* := *T* × 0.*9995* for every 4*S ×* 10^™3^ iterations, where *S* ≥ 5 *×* 10^4^ is the total number of iterations. RJMCMC samples an alternative model *M*_new_ from a proposed distribution *P*^prop^(*M*_new_|*M*). The new model *M*_new_ is sampled from proposals for the birth and death of particles, moving and rotating of particles, connections and reconnections between particles, and the track-birth and track-death of particles. The RJMCMC sampling starts with the empty model *M*_*0*_ = {*𝛷, 𝛷}* without any particles or edges, and at each iteration, one of the proposals is randomly chosen. Whether *M*_new_ is accepted is determined by ratio *R*, as follows:

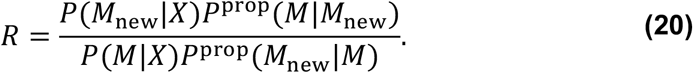

If *R* ≥ 1, *M*_new_ will be accepted; if *R <* 1, *M*_new_ will be accepted with probability *R*. By substituting Eq. 7 and 9 into Eq. 20, to obtain the following:

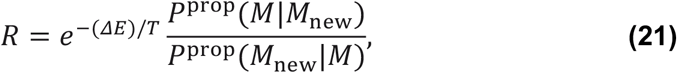

where *𝛥E* = *E*{*X*}(*M*_new_) − *E*{*X*}(*M*) and *E*{*X*}(*⋅*) = *E*_ext_(*⋅, X*) + *E*_int_(*⋅*) is the total energy function of a model that represents the F-actin network in HS-AFM image *X*.

### Details of the proposals

A new model was proposed by reorganizing the connected structures of the particles in the current model with the MCMC sampling. The design of proper proposals is important to help prevent their rejection; proposals with completely random reorganization of the model must be unrealistic and cannot fit the AFM images, and consequently the rejection ratio is high. Biologically realistic sampling was thus adopted to efficiently determine the optimal model that maximizes or nearly maximizes *P*(*M*|*X*). In this study, six proposal types were uniformly sampled: (1) particle birth; (2) particle death; (3) moving and/or rotating of particles; (4) connection/reconnection of edges; (5) track-birth of particle; and (6) track-death of particle.

#### Birth proposal

The birth proposal generates a new particle *p*_new_ in the current model, and the position of 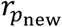 was sampled from the distribution as follows:

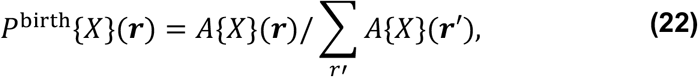

where *A{X*}(***r***) = m*ax(*m*ax*_*n*_ *Z*′*{X*}(***r, n***),0) + *ϵ*. The small constant *ϵ* = 0.001 ensures that particles can appear at any location in the HS-AFM image *X. Z*′*{X*}(***r, n***) is the normalized AOD function, which is defined as follows:

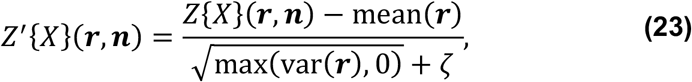

where *𝜁* = 0.01 is a small constant, and m*ean*(***r***) and *var(****r***) represent the local average and variance of the AOD function around image position ***r***, respectively, as follows:

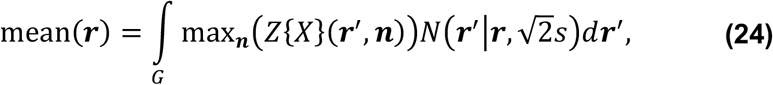

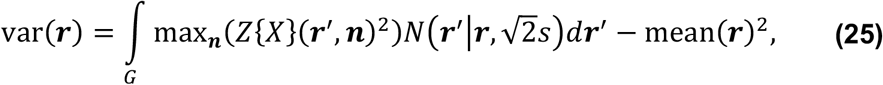

where *N*(. |*μ, σ*) indicates a normal distribution with mean *μ* and variance *σ*^*2*^, and *s* indicates a constant set to be the scale of the particles, i.e., *s* = 2. The orientation of *p*_new_ is given by 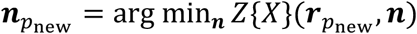 for efficient realistic proposals, instead of the random sampling previously described^19,21^. After generating a new particle for the model, the change in energy was found to be as follows; *𝛥E* = *C*_coll_(*p*_new_) + *E*_ext_{*X*}(*p*_new_). The acceptance ratio of the birth proposal is:

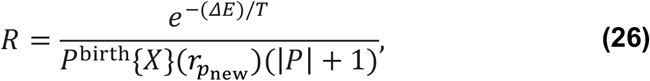

where |*P*| means the total number of particles in the model.

#### Death proposal

The death proposal randomly selects an existing particle *p*_*0*_ for removal from the current model. If the selected particle *p*_*0*_ is connected to others, the death proposal is rejected. Otherwise, *p*_*0*_ is removed using the acceptance ratio:

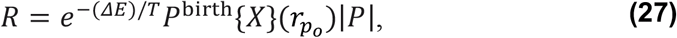

where *𝛥E* = *C*_coll_(*p*_*0*_) + *E*_ext_{*X*}(*p*_*0*_).

#### Moving and/or rotating proposal

The moving and rotating proposals randomly select particle *p* from the current model *M* and then randomly determine whether to propose altering its position ***r***_*p*_ and orientation ***n***_*p*_. We denote *p*′ as the particle selected after an attribute change is proposed. If the rotating proposal is selected, *n*_*p’*_ is updated by the normalized result of ***n***_*p*_ + ***ϵ***_*n*_, where ***ϵ***_*n*_ is the Gaussian noise with zero mean and covariance matrix *diag*(*ϵ*_*n*_, *ϵ*_*n*_) and *ϵ*_*n*_ is chosen to be 1.5 empirically. If the moving proposal is selected, then ***r***_*p’*_ = ***r***_*p*_ + ***ϵ***_*r*_, where ***ϵ***_*r*_ is the Gaussian noise with zero mean and covariance matrix *diag*(*ϵ*_*r*_, *ϵ*_*r*_), and *ϵ*_*r*_ is chosen to be 2 empirically. If the proposal is to change both, the position and orientation of the selected particle will be changed simultaneously, as described above. These proposals for *p*′ are accepted at

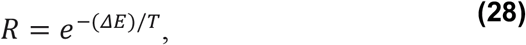

where

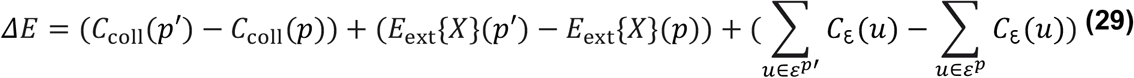

and *ε*^*p*^ is the set of all edges connected to a particle *p*.

#### Connection/reconnection proposal

*T*he connection/reconnection proposal either reconnects the existing edges or adds new edges to the terminal particles in the model. The proposal uniformly samples a connecting site 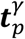 (black point in **Fig. 11**) of particle *p* from the current model. If there is a *u*_*0*_ edge at 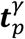, the proposal reconnects the edge as follows; first *u*_*0*_ is removed and then a set of candidate edges is created in the vicinity as follows:

**Figure 11:**
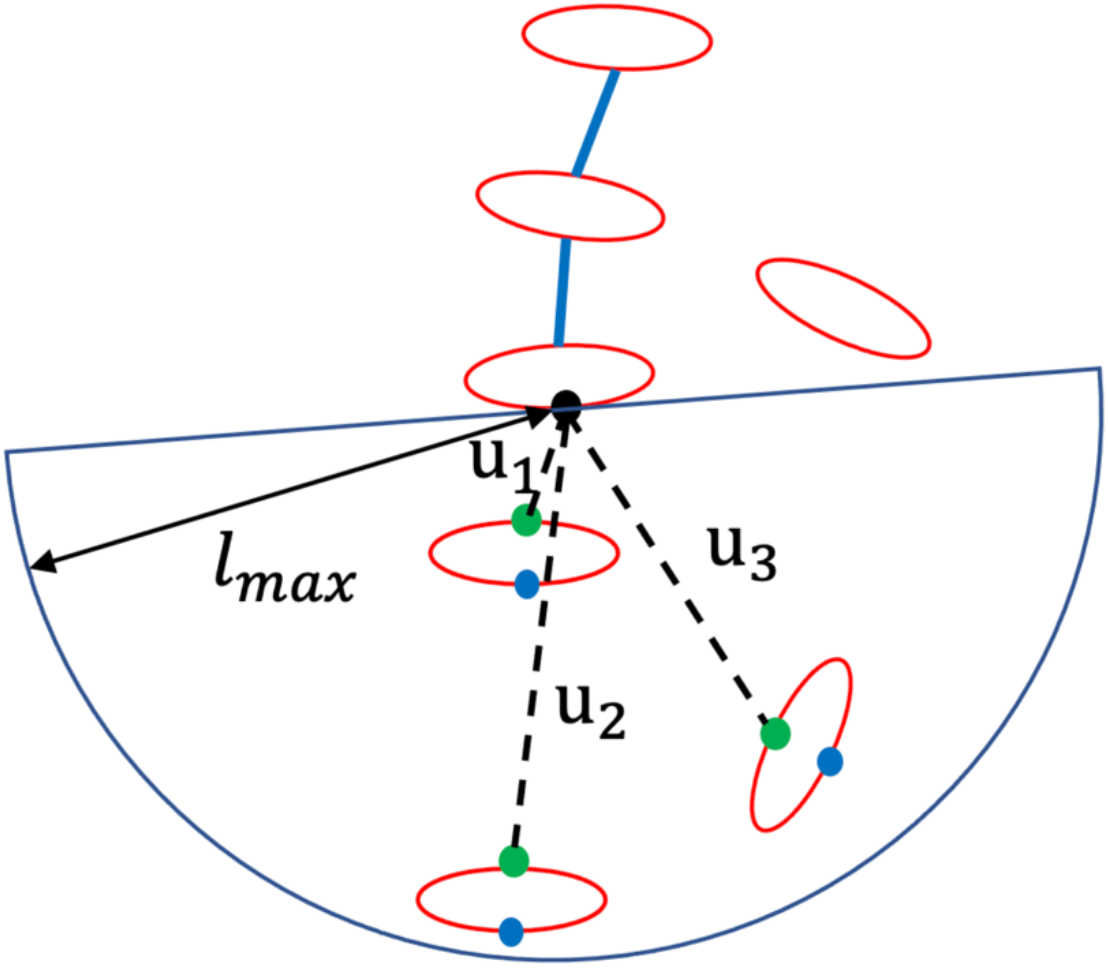
Creation of candidate edges in the connection/reconnection proposal. Connection or reconnection sites were randomly selected (black points). Candidate edges *ε*^cand^ were searched for within a semicircle area with a radius of *l*_max_, which was generated from the end particle toward its orientation. The free connecting sites facing the selected connecting site (green points) are candidates, whereas those facing the opposite direction (blue points) are ignored. The candidate edge with minimal elastic energy was selected as the new edge for MCMC sampling. In this case, *u*_1_ is selected.

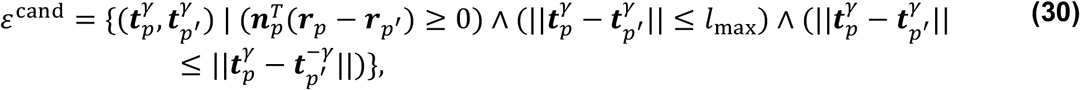

where *p*′ indicates candidate particles for reconnection, 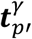 is the connecting site of *p*′ that no edge is connecting to, and *l*_*ki*5_ = 4 is the searching radius. In contrast to that in previous studies^19,21^, to increase the efficiency of the algorithm, the conditions 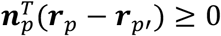 and 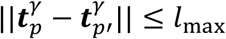 were used to ensure that the set of candidate edges is located in front of 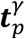 with the searching radius *l*_max_ (blue semicircle in **Fig. 11**). The condition 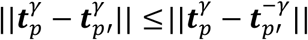 ensures that the connecting site 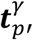 of the candidate particle *p*′ inside the searching area is chosen to be the one that faces 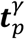 (green points in **Fig. 11**). The new edge *u*′_*0*_ is selected to have minimal elastic energy in *ε*^cand^, as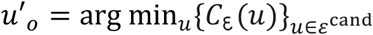, instead of the random sample method used in previous studies^19,21^. The proposal of *u*′_*0*_ is accepted as:

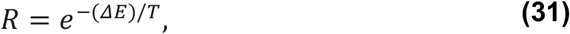

where *𝛥E* = *C*_દ_(*u*′_*0*_) − *C*_દ_(*u*_*0*_).

If the selected connecting site has no edges, the proposed method adds an edge from the set of candidate edges in Eq. 30 and follows the same method as described above. In this case, *𝛥E* = *C*_દ_(*u*′_*o*_).

#### Track-birth proposal

The track-birth proposal adds a new particle and connects it to a terminal particle with an edge. The proposal randomly selects a particle *p* from *P*_*term*_, which is the set of terminal particles *P*_*term*_ *⊆ P* in model *M*. The unconnecting site for the selected terminal particle is 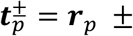 *w**n***_*p*_ The position for adding particle *p*_new_ is sampled from a Gaussian distribution as 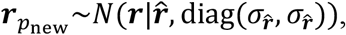 where 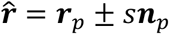, and 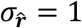 was determined empirically. This sampling ensures that *p*_new_ is located as if the existing F-actin were elongated in a straight line. Note that the orientation of *p*_new_ is given by 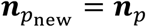 instead of sampling from a distribution^19,21^ to facilitate efficient sampling. The new edge connecting *p* and *p*_new_ *i*s created as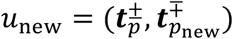. The acceptance ratio is:

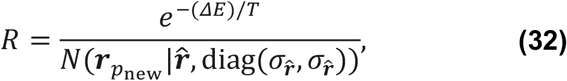

where *𝛥E* = *C*_coll_(*p*_new_) + *E*_ext_{*X*}(*p*_new_) + *C*_દ_(*u*_new_).

#### Track-death proposal

The track-death proposal randomly selects a terminal particle *p*_*0*_ from *P*_*term*_ for removal. If the second particle from the terminal *p*_*0*_ (denoted by *p*) does not have two edges, the track-death proposal is rejected, otherwise, *p*_*0*_ is removed using the acceptance ratio:

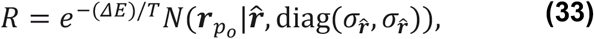

where *𝛥E* = −(*C*_coll_(*p*_*0*_) + *E*_ext_{*X*}(*p*_*0*_) + *C*_દ_(*u*_*0*_)), and *u*_*0*_ indicates the edge connecting *p*_*0*_ and *p*.

## Code availability

The source code will be released after the paper is accepted.

## Data availability

No datasets were generated or analyzed during the current study.

## Acknowledgments

The authors thank Mr. N. Sakai and Mr. Y. Itagaki for their technical assistance with the HS-AFM imaging. This work was supported by JSPS KAKENHI (grant numbers JP18H02436, JP18KK0196, and JP22H05171; to S.H.Y.), and the Japan Agency for Medical Research and Development (AMED; grant number JP18gm5810018; to S.H.Y.). This study was also supported in part by the Grant-in-Aid for Challenging Research (Exploratory; grant number 19K22422 to H.N. and S.H.Y.] from the Japan Society for the Promotion of Science (JSPS), the Moonshot R&D–MILLENNIA Program (grant number JPMJMS2024-9; to J.S.T.), Grant-in-Aid for Transformative Research Areas (B) (grant number 21H05170), and the Cooperative Study Program of Exploratory Research Center on Life and Living Systems (ExCELLS; program number 19-102 to H.N.).

## Author Contributions

H.N. and S.H.Y. conceived the study. H.J., H. S., and H.N. developed the method; H.J. implemented the method and performed mathematical and data analyses; M. F., H.J., and H.N. proposed the F-actin organization mechanism in the cell cortex. All authors contributed to the writing of the manuscript.

## References

1. Blanchoin, L., Boujemaa-Paterski, R., Sykes, C., and Plastino, J. (2014). Actin Dynamics, Architecture, and Mechanics in Cell Motility. Physiol Rev 94, 235–263. 10.1152/physrev.00018.2013.

2. Nonaka, S., Naoki, H., and Ishii, S. (2011). A multiphysical model of cell migration integrating reaction–diffusion, membrane and cytoskeleton. Neural Networks 24, 979–989. 10.1016/j.neunet.2011.06.009.

3. Mullins, R.D., Heuser, J.A., and Pollard, T.D. (1998). The interaction of Arp2/3 complex with actin: Nucleation, high affinity pointed end capping, and formation of branching networks of filaments. Proceedings of the National Academy of Sciences 95, 6181–6186. 10.1073/pnas.95.11.6181.

4. Yamao, M., Naoki, H., Kunida, K., Aoki, K., Matsuda, M., and Ishii, S. (2015). Distinct predictive performance of Rac1 and Cdc42 in cell migration. Sci Rep 5, 17527. 10.1038/srep17527.

5. Imoto, D., Saito, N., Nakajima, A., Honda, G., Ishida, M., Sugita, T., Ishihara, S., Katagiri, K., Okimura, C., Iwadate, Y., et al. (2021). Comparative mapping of crawling-cell morphodynamics in deep learning-based feature space. PLoS Comput Biol 17, e1009237. 10.1371/journal.pcbi.1009237.

6. Vinzenz, M., Nemethova, M., Schur, F., Mueller, J., Narita, A., Urban, E., Winkler, C., Schmeiser, C., Koestler, S.A., Rottner, K., et al. (2012). Actin branching in the initiation and maintenance of lamellipodia. J Cell Sci. 10.1242/jcs.107623.

7. Maly, I. V., and Borisy, G.G. (2001). Self-organization of a propulsive actin network as an evolutionary process. Proceedings of the National Academy of Sciences 98, 11324–11329. 10.1073/pnas.181338798.

8. Mullins, R.D., Stafford, W.F., and Pollard, T.D. (1997). Structure, Subunit Topology, and Actin-binding Activity of the Arp2/3 Complex from Acanthamoeba. Journal of Cell Biology 136, 331–343. 10.1083/jcb.136.2.331.

9. Vignjevic, D., Yarar, D., Welch, M.D., Peloquin, J., Svitkina, T., and Borisy, G.G. (2003). Formation of filopodia-like bundles in vitro from a dendritic network. Journal of Cell Biology 160, 951–962. 10.1083/jcb.200208059.

10. Yang, C., and Svitkina, T. (2011). Visualizing branched actin filaments in lamellipodia by electron tomography. Nat Cell Biol 13, 1012–1013. 10.1038/ncb2321.

11. Yang, C., and Svitkina, T. (2011). Filopodia initiation. Cell Adh Migr 5, 402–408. 10.4161/cam.5.5.16971.

12. Winkler, C., Vinzenz, M., Small, J.V., and Schmeiser, C. (2012). Actin filament tracking in electron tomograms of negatively stained lamellipodia using the localized radon transform. J Struct Biol 178, 19–28. 10.1016/j.jsb.2012.02.011.

13. Suzuki, Y., Sakai, N., Yoshida, A., Uekusa, Y., Yagi, A., Imaoka, Y., Ito, S., Karaki, K., and Takeyasu, K. (2013). High-speed atomic force microscopy combined with inverted optical microscopy for studying cellular events. Sci Rep 3, 2131. 10.1038/srep02131.

14. Yoshida, A., Sakai, N., Uekusa, Y., Deguchi, K., Gilmore, J.L., Kumeta, M., Ito, S., and Takeyasu, K. (2015). Probing in vivo dynamics of mitochondria and cortical actin networks using high-speed atomic force/fluorescence microscopy. Genes to Cells 20, 85–94. 10.1111/gtc.12204.

15. Zhang, Y., Yoshida, A., Sakai, N., Uekusa, Y., Kumeta, M., and Yoshimura, S.H. (2017). In vivo dynamics of the cortical actin network revealed by fast-scanning atomic force microscopy. Microscopy 66, 272–282. 10.1093/jmicro/dfx015.

16. Yamao, M., Aoki, K., Yukinawa, N., Ishii, S., Matsuda, M., and Naoki, H. (2016). Two New FRET Imaging Measures: Linearly Proportional to and Highly Contrasting the Fraction of Active Molecules. PLoS One 11, e0164254. 10.1371/journal.pone.0164254.

17. Green, P.J. (1995). Reversible jump Markov chain Monte Carlo computation and Bayesian model determination. Biometrika 82, 711–732. 10.1093/biomet/82.4.711.

18. Reisert, M., and Kiselev, V.G. (2011). Fiber Continuity: An Anisotropic Prior for ODF Estimation. IEEE Trans Med Imaging 30, 1274–1283. 10.1109/TMI.2011.2112769.

19. Skibbe, H., Reisert, M., Nakae, K., Watakabe, A., Hata, J., Mizukami, H., Okano, H., Yamamori, T., and Ishii, S. (2019). PAT—Probabilistic Axon Tracking for Densely Labeled Neurons in Large 3-D Micrographs. IEEE Trans Med Imaging 38, 69–78. 10.1109/TMI.2018.2855736.

20. Reisert, M., and Skibbe, H. (2011). Steerable Deconvolution Feature Detection as an Inverse Problem. In, pp. 326–335. 10.1007/978-3-642-23123-0_33.

21. Skibbe, H., Reisert, M., Maeda, S., Koyama, M., Oba, S., Ito, K., and Ishii, S. (2015). Efficient Monte Carlo Image Analysis for the Location of Vascular Entity. IEEE Trans Med Imaging 34, 628–643. 10.1109/TMI.2014.2364404.

22. Ott, A., Magnasco, M., Simon, A., and Libchaber, A. (1993). Measurement of the persistence length of polymerized actin using fluorescence microscopy. Phys Rev E 48, R1642–R1645. 10.1103/PhysRevE.48.R1642.

23. Svitkina, T.M. (2020). Actin Cell Cortex: Structure and Molecular Organization. Trends Cell Biol 30, 556–565. 10.1016/j.tcb.2020.03.005.

24. Chugh, P., Clark, A.G., Smith, M.B., Cassani, D.A.D., Dierkes, K., Ragab, A., Roux, P.P., Charras, G., Salbreux, G., and Paluch, E.K. (2017). Actin cortex architecture regulates cell surface tension. Nat Cell Biol 19, 689–697. 10.1038/ncb3525.

25. Nakamura, F., Osborn, T.M., Hartemink, C.A., Hartwig, J.H., and Stossel, T.P. (2007). Structural basis of filamin A functions. Journal of Cell Biology 179, 1011–1025. 10.1083/jcb.200707073.

26. Weichsel, J., Urban, E., Small, J.V., and Schwarz, U.S. (2012). Reconstructing the orientation distribution of actin filaments in the lamellipodium of migrating keratocytes from electron microscopy tomography data. Cytometry Part A 81A, 496–507. 10.1002/cyto.a.22050.

27. Aguet, F., Jacob, M., and Unser, M. (2005). Three-dimensional feature detection using optimal steerable filters. In IEEE International Conference on Image Processing 2005 (IEEE), pp. II–1158. 10.1109/ICIP.2005.1530266.

28. Reisert, M., and Burkhardt, H. (2008). Complex Derivative Filters. IEEE Transactions on Image Processing 17, 2265–2274. 10.1109/TIP.2008.2006601.

